# A Meta-Analysis of Functional Neuroimaging Tasks associated with Perceptual Pseudoneglect

**DOI:** 10.1101/2025.02.04.636501

**Authors:** Mura Abdul-Nabi, Matthias Niemeier

## Abstract

Major evidence for a right-hemisphere dominance of the brain in spatial and/or attentional tasks comes from lesion studies in patients with spatial neglect. However, the neuroanatomy of the different forms of neglect remains a matter of debate, and it remains unclear how dysfunctions in neglect relate to intact processes. In the healthy brain, perceptual pseudoneglect has been considered to be a phenomenon complementary to specific subtypes of neglect as observed in paradigms such as the line bisection task. Therefore, the current study investigated the intact functional anatomy of perceptual pseudoneglect using a meta-analysis to compensate for some of the limitations of individual imaging studies. We collated the data from 24 articles that tested 952 participants with a range of paradigms (landmark task, line bisection, grating-scales task, and number line task) obtaining 337 foci. Using Activation Likelihood Estimation (ALE) we identified a right-hemisphere biased network of cortical areas, including superior and intraparietal regions, the intraoccipital sulcus together with other occipital regions, as well as inferior frontal areas that were associated with perceptual pseudoneglect in partial agreement with lesion studies in patients with neglect. Our study is the first meta-analysis on the mechanisms underlying perceptual judgments which have been shown to give rise to perceptual pseudoneglect.

## Introduction

Spatial neglect is a set of spatial cognitive deficits that have been summarized as inabilities to report, respond, or orient to stimuli contralateral to the side of a brain damage (e.g., Heilman et al., 1993). Neglect is more commonly observed after right than left brain lesions, and thus it is an important indicator for a right-hemisphere dominance of the neural mechanisms underlying visual attention and spatial awareness. As such neglect has garnered significant interest in the literature. Yet, several questions remain unresolved. Crucially, it is unclear how neglect symptoms map onto brain damage.

A recent meta-analysis of lesion studies in neglect patients collated a list of best-practice criteria for lesion-symptom mapping (Moore et al., 2023; also see Bates et al., 2003; de Haan and Karnath, 2018; Sperber and Karnath, 2017; Thiebaut de Schotten et al., 2014) showing none of the relevant studies attained a perfect score. Also, depending on the score, different lesion patterns emerged with varying core lesions in the supramarginal gyrus, the superior longitudinal fasciculus, the insular cortex, or the postcentral gyrus. Furthermore, the analysis found substantial differences in functional anatomy between different subtypes of neglect. Also consistent with the heterogeneity of neglect, lesion patterns have been found to vary as a function of diagnostic tests (Molenberghs et al., 2012).

To complicate matters functional deficits and brain plasticity after brain lesions interact in a complex and difficult-to-observe manner (Pascual-Leone et al., 2005). For example, extinction, a deficit akin to neglect that affects contralesional stimuli presented together with competing ipsilesional stimuli, does not occur immediately after simulated brain lesions induced by inhibitory continuous theta-burst stimulation (Vesia et al., 2015). Instead, the authors found extinction to arise several minutes after stimulation, arguably resulting from short-term plasticity due to interactions between the two hemispheres. Interhemispheric interactions also seem to play an essential role in neglect in general (Koch et al., 2008; also see Corbetta & Shulman, 2011) as well as in related asymmetries in the intact brain (Le et al., 2015). This suggests that behavioural impairments and the site of damage do not necessarily relate to one another in a straight-forward manner, contrary to one of the central assumptions of lesion studies (Caramazza, 1984).

Given these inherent challenges of lesion studies, a comprehensive picture of the functional anatomy of neglect calls for comparisons between research in patients with imaging studies in healthy participants to estimate to which extent lesion patterns in neglect agree with the anatomy of intact processes that can be assumed to be affected in spatial neglect. One such set of functions involves perceptual judgements about visuo-spatial features of horizontal lines or bars such as length (e.g., Harvey & Milner, 1995), luminance (Mattingley et al., 1994), or spatial frequency (Niemeier et al., 2007). These tasks now have been validated to yield comparable visual field differences in the intact brain called perceptual pseudoneglect (Chen et al., 2019; see a detailed discussion of construct validity there) or attentional bias (Bultitude et al., 2006; McCourt et al., 2005; Singh et al., 2011). Furthermore, the differences seem to reflect approximately complementary, although much less severe, asymmetries compared to deficits found in neglect patients (Bowers & Heilman, 1980; McCourt & Jewell, 1999) when performing line length judgements in the manual line bisection task that is commonly used in clinical assessments (e.g., Jewell & McCourt, 2000).

Line bisection and other perceptual judgment tasks have been employed in several imaging studies with a variety of designs. However, most of these studies used simple contrasts with only one experimental and one control condition (see Table 1) thereby bearing a greater risk of false positive and false negative errors in the identification of relevant brain areas. This is particularly an issue for perceptual judgment paradigms given that they permit test participants to use a variety of tasks strategies.

**Table 1.** Studies included in the meta-analysis.

| Publication | N | Contrast | Foci | Analysis |
| --- | --- | --- | --- | --- |
| Badzakova-Trajkov et al. (2010) | 84 | LM > Bisector Detection | 7 | Omnibus; Landmark |
| Badzakova-Trajkov et al. (2011) | 199 | LM > Bisector Detection | 9 | Omnibus; Landmark |
| Cai et al. (2013) | 31 | LM > Bisector Detection<br>(includes L and R language dominance) | 19 | Omnibus; Landmark |
| Cavezian et al. (2011) | 9 | LM w/ greyscales > Bisector Detection | 27 | Omnibus; Landmark |
|  |  | LB w/ greyscales > Saccade Task | 14 | Omnibus; Line Bisection |
| Chen et al. (2020) | 21 | Latent variable modelling a continuum of the grating-scales task/high spatial frequency vs. low spatial frequency vs. frame condition vs. fixation | 21 | Omnibus |
| Cicek et al. (2009) | 11 | LM > Bisector Detection | 6 | Omnibus; Landmark |
|  |  | LB > Move cursor to edge | 3 | Omnibus; Line Bisection |
| Fink et al. (2000a) | 12 | LM > Bisector Detection | 7 | Omnibus; Landmark |
| Fink et al. (2000b) | 12 | LM > Bisector Detection | 5 | Omnibus; Landmark |
| Fink et al. (2001) | 11 | LMV + LM > Bisector Detection | 13 | Omnibus |
| Fink et al. (2002) | 12 | LM + LL > Instructions | 12 | Omnibus |
| Fink et al. (2003) | 12 | LM (eye movements) > Bisector Detection | 9 | Omnibus; Landmark |
| Flöel et al. (2005) | 7 | LM > Overlapping waveform line detection | 26 | Omnibus; Landmark |
| Hurwitz et al. (2011) | 16 | DB > Saccade task | 3 | Omnibus; Line Bisection |
| Liu et al. (2019) | 34 | Number LB > Number Recognition Task | 37 | Omnibus; Number Line |
|  |  | LB > Dot Position Recognition Task | 11 | Omnibus; Line Bisection |
| Marshall et al. (1997)* | 9 | LM > Line Flicker Task | 1 | Omnibus; Landmark |
| Plewan et al. (2012) | 24 | LM Mueller Lyer > Rest | 5 | Omnibus; Landmark |
| Revill et al. (2011) | 26 | LM > Rest | 15 | Omnibus; Landmark |
|  |  | LM > search task | 13 | Omnibus |
| Sahan et al. (2019) | 24 | Number LB > Shape Recognition Task (Encoding & Decision-Making Phases) | 26 | Omnibus; Number Line |
| Schuster et al. (2017) | 15 | LM (Version C) > Bisector Detection (Session 1 & 2) | 31 | Omnibus; Landmark |
| Weidner et al. (2007) | 15 | LM Mueller Lyer > Luminance Detection | 2 | Omnibus; Landmark |
| Weiss et al. (2000) | 12 | Near + Far LB > Pointing Task | 6 | Omnibus; Line Bisection |
| Weiss et al. (2003) | 12 | Near + Far LM > Near + Far LB | 16 | Omnibus |
|  |  | Near + Far LB > Near + Far LM | 7 | Omnibus; Line Bisection |
| Zago et al. (2016) | 293 | LM > Saccade Task | 11 | Omnibus; Landmark |
| Zago et al. (2017) | 51 | LM > Saccade Task | 10 | Omnibus; Landmark |

LM: Landmark task; LB: Line Bisection task; LMV: Vertical Landmark Task; LL: Line Length
Task; DB: Dynamic Bisection Task. \*The study employed SPECT.

A better approach is to design a control condition with physically identical stimuli that are used with different task instructions (e.g., Revill et al., 2011). However, as a potential limitation of this approach, it is possible that brain areas that are critical for some function or task A, might nevertheless yield a stronger BOLD signal in some other task B. This way, subtracting task B activity from task A activity would potentially remove an essential area from the results. For example, Filimon and colleagues (2009) have found that the frontal eye fields activate more during reach tasks than eye movement tasks. Thus, subtracting BOLD signal associated with reaching from the BOLD signal associated with eye movements could obscure eye movement-related activation in the frontal eye fields even though they are critically involved in eye movements.

A third approach then is to use more complex experimental designs for better experimental control. Benwell et al. (2014) used a factorial design for an EEG experiment with landmark tasks with long and short lines and with a control task that required bisections to be detected. To our knowledge a factorial design has not been used in an fMRI experiment on the attentional bias. Chen et al. (2020) used partial-least squares to model a cognitive vector based on four tasks that were incrementally more likely to involve brain regions to do with perceptual judgments that yield pseudoneglect biases. However, not all perceptual judgment paradigms lend themselves to such a set of tasks. The arguably simplest way to attain better control would be a conjunction analysis where multiple contrasts of experimental and control tasks are looked at together such that confounds of one contrast are compensated for by other contrasts (Price & Friston, 1997). To our knowledge to date no pseudoneglect study has used conjunction analysis.

However, it can be assumed that a meta-analysis, although different from a conjunction analysis, collates multiple contrasts from potentially different paradigms (depending on the paradigms of the included studies) and this way potentially discards some of the suspected false positive and false negative errors of the included studies. Therefore, the aim of the present study was to conduct a meta-analysis of perceptual judgments tasks that are known to yield perceptual pseudoneglect to improve overall validity. Furthermore, the question was whether the resulting functional neuroanatomy would be mirrored in patients exhibiting line bisection deficits after lesions along the intraparietal and intraoccipital sulci (IPS and IOS), medial occipital cortex and anterior portions of the middle frontal gyrus (MFG; see meta-analysis in Molenberghs et al., 2012).

## Materials and Methods

To ensure a comprehensive article list was obtained, imaging studies using perceptual judgments to measure attentional bias / pseudoneglect published on or prior to March 10, 2025 were obtained by searching the PubMed (https://pubmed.ncbi.nlm.nih.gov/), PsycInfo (https://www.apa.org/pubs/databases/psycinfo), and Web of Science (https://www.webofknowledge.com/) databases using the following keyword search: “(line bisection OR landmark task OR pseudoneglect OR greyscales task OR grayscales task OR grating-scales task) AND (fMRI OR PET).” Reference lists were also checked to avoid missing relevant articles. A repetition of the search on November 29, 2023, yielded no additional relevant publications.

Articles were selected according to the following inclusion criteria: 1) studies were published in English, 2) participants were healthy, neurotypical adults (18-65 years old), 3) studies assessed attentional biases using either horizontal line bisection, landmark, greyscales, or gratingscales tasks, 4) fMRI or PET was used to assess neural activity (one study using SPECT was included as well; additional searches revealed that no other relevant study used this method), and 5) Montreal Neurological Institute (MNI) or Talairach standardized coordinates of activation were provided. The exclusion criteria were as follows: 1) participants with a history of brain damage or other clinical populations, 2) clinical or drug related interventions, 3) studies on children (< 18 years old) or seniors (65+ years old), and 4) meta-analyses or literature reviews as evidenced based on inspecting the articles’ titles or content. Additionally, we only included those contrasts that compared task activation to a control or baseline condition and that imaged the entire brain. Using these criteria, we selected 24 articles consisting of 952 subjects and 337 foci of activation to be included in this meta-analysis. An inter-rater agreement of r = 0.93 was obtained between the authors’ selections of studies to be included. Any discrepancy was then resolved through closer inspection of the articles in question. Figure 1 details our literature search and selection process.

**Figure 1.**
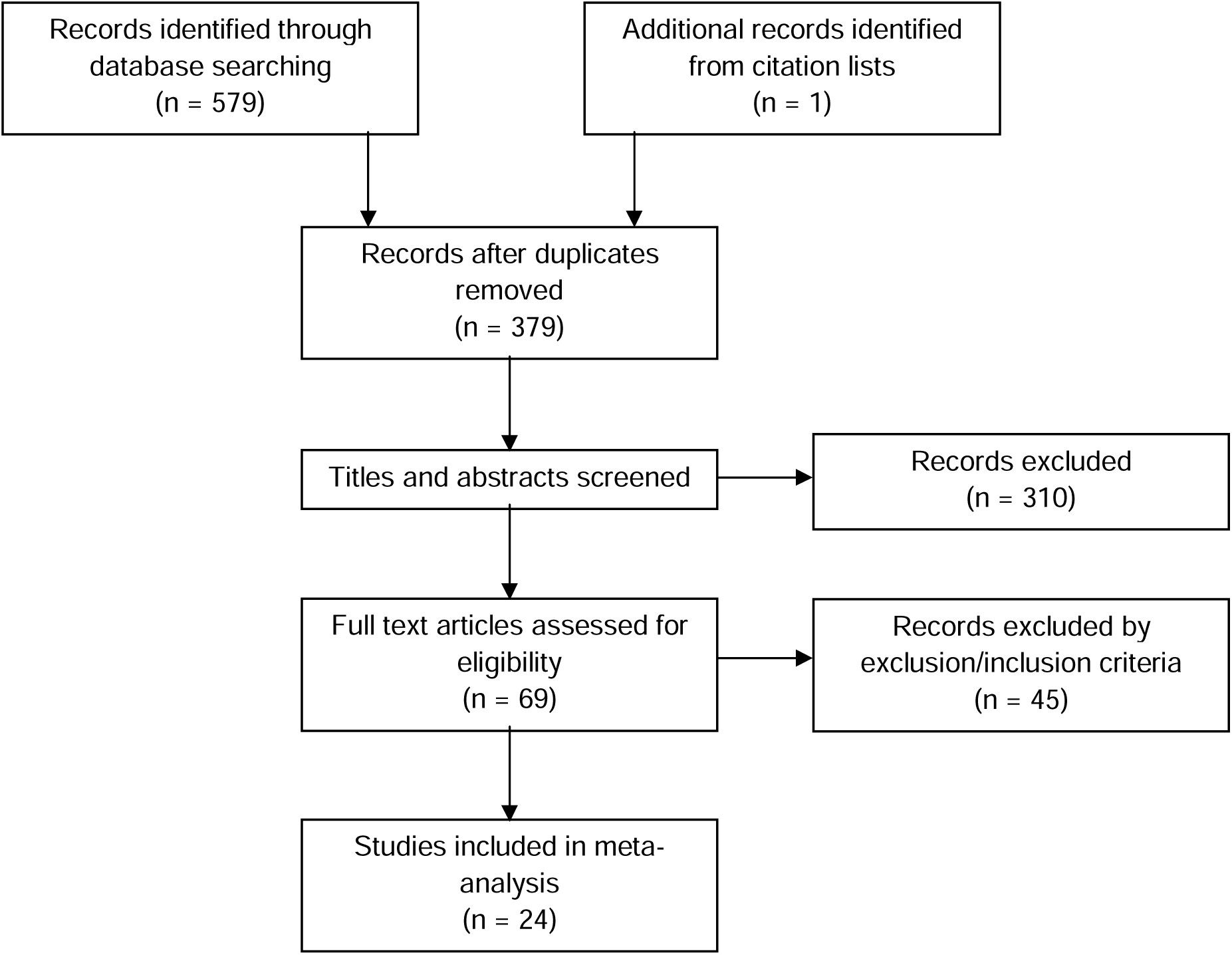
Outline of the literature search process.

Upon finalizing the list of studies, we converted all coordinates of activation reported in Talairach standardized space to MNI space using the mni2tal transform (https://bioimagesuiteweb.github.io/webapp/mni2tal.html). These data were then submitted to an omnibus meta-analysis (24 articles consisting of 952 subjects and 337 foci). Next, we categorized studies based on the paradigms used to assess pseudoneglect to allow for comparison among tasks. These included: landmark tasks, physical line bisection tasks, and number line bisection tasks. The meta-analysis for the Landmark task included 16 studies involving 810 subjects and 173 foci. Notably, the majority of these studies did not report behavioural data on perceptual biases. We decided to include all studies (rather than introduce a post-hoc inclusion criterion) as our objective was to investigate the activation patterns associated with perceptual judgment tasks previously shown to elicit perceptual pseudoneglect (e.g., Chen et al., 2019). Restricting the analysis to studies that reported perceptual biases would have introduced a selection bias, potentially favouring studies that tested more strongly lateralized participants and thereby skewing the functional data towards exaggerated brain asymmetries. Here, we selected contrasts that employed baseline/control conditions. (No individual meta-analyses were performed for the Line Bisection task, 6 studies, 94 subjects, 44 foci, the Number Line task, 2 studies, 58 subjects, 52 foci, and the grating-scales task, 1 study, 21 subjects, 21 foci). Multiple contrasts from a single experiment were pooled together to avoid greater weight being given to multiple analyses with the same set of participants (Müller et al., 2018). A complete list of studies and their attributes can be found in Table 1.

For each meta-analysis, data were analyzed using the Activation Likelihood Estimation (ALE) method outlined in Brainmap GingerALE Version 3 (http://brainmap.org/ale/). The algorithm accounts for spatial uncertainty by treating individual foci as centers of 3D Gaussian probability distributions with a width that depends on the sample size of the respective study (i.e., the width is inversely related to the square root of the sample size). Summing over the Gaussians then results in a probability distribution that is superimposed with a grey matter mask. Next, a permutation procedure computes nonparametrically a null-distribution of random spatial association between experiments to identify statistically significant convergence of activation foci. Here, we employed a cluster level approach with an uncorrected P value of p < 0.001 and a cluster-level family-wise error (FWE) set at p < 0.05 for the omnibus analysis. An additional meta-analysis for the landmark data alone was slightly below the number of studies recommended for ALE (Eickhoff et al., 2016). To avoid unstable results we chose a more conservative p-value of 0.001 for the cluster-level FWE. Threshold permutations were set at 1000, as per recommendations outlined in Eickhoff et al. (2012). Results of the ALE analyses were visualized using MRIcron (http://www.cabiatl.com/mricro/mricron/) and BrainNet Viewer (https://www.nitrc.org/projects/bnv/; Xia et al., 2013). BrainNet Viewer was also used to visualize activation foci for the line bisection (6 studies) and the number line task (2 studies) and plotted together with the activation maps for grating-scales task (Chen et al., 2020). Given the small numbers of these studies only qualitative syntheses and comparisons instead of meta-analyses were used.

## Results

### The functional anatomy of the perceptual attentional bias: omnibus analysis

Figure 2 shows the results of the omnibus meta-analysis, including all 24 studies. Significant overlap in activation was found in posterior regions, including the anterior and posterior intraparietal sulcus (IPS) that was more prominent in the right than the left hemisphere and reached into the superior parietal lobule (SPL). Exclusively right hemisphere overlap was found in the intraoccipital sulcus (IOS). Likewise, exclusive right-brain overlap was found in frontal regions for the precentral sulcus at the intersection with the superior frontal sulcus, probably the frontal eye field (FEF) and at the level of the inferior frontal cortex (IFC). In addition, there was right dorsolateral prefrontal cortex (DLPFC) activity and dorso-medial activity near the pre-supplementary motor area. Overlap in the anterior insula (aIns) was bilateral. Notably, several studies reported foci in the cerebellum but there was no significant overlap. We confirmed these observations statistically using the AAL3 atlas (Rolls et al., 2020) to create independently defined masks. In terms of posterior regions, as defined by the atlas, we found significantly more (or exclusive) right-brain involvement for the SPL, IPS (i.e., the inferior parietal lobule, not including supramarginal and angular gyrus), as well as the superior and middle occipital gyrus (t’s≥10.85, p<0.001) with spurious involvement of the right but not left postcentral and angular gyrus (t’s≥3.84, p’s≤0.001). As for AAL3-defined frontal regions we found exclusive right hemisphere involvement of the superior frontal gyrus, middle frontal gyrus, and precentral gyrus (t’s≥4.27, p’s≤0.001) and right-dominant involvement for the SMA as well as the opercular and (spuriously) triangular part of the inferior frontal gyrus, both of which were reaching into the aIns (t’s≥4.50, p’s≤0.001). The remainder of the insula showed no significant left-vs. right-side difference (t=0.77, p=0.441).

**Figure 2.**
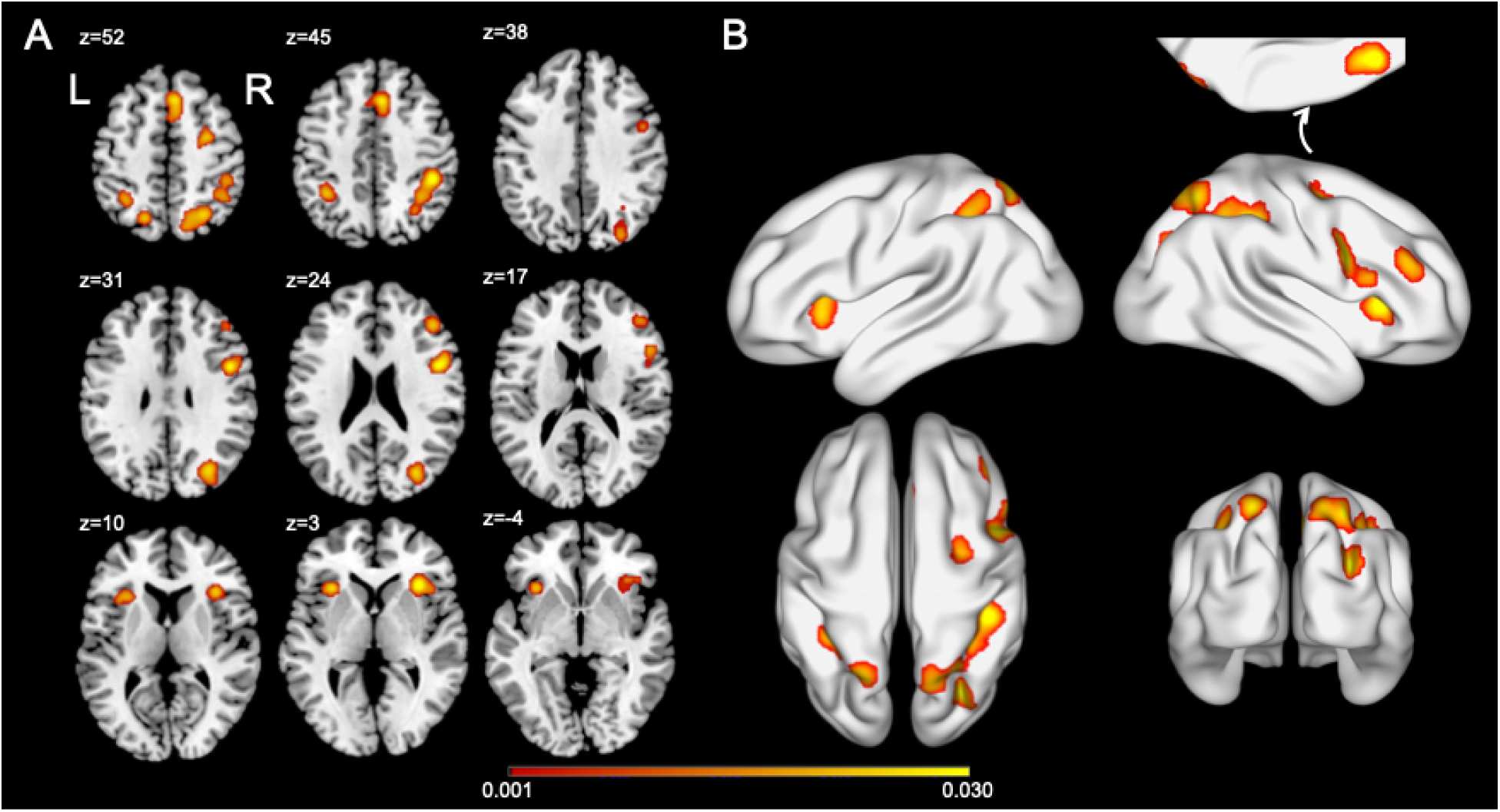
Results of the omnibus meta-analysis. A. Horizontal slices. B. Lateral, dorsal, and posterior views. The colour bar represents ALE scores found to be significant for a cluster-level family-wise error (FWE) set at p < 0.05.

### Influence of task on the cortical network of the perceptual attentional bias

Perceptual forms of attentional biases have been captured with several different kinds of tasks, although not all these tasks have been frequently used in imaging studies such that more commonly used paradigms have a stronger influence on our omnibus results. To examine the effect of task on brain activation we inspected the data for specific tasks, respectively. Only for the landmark task there were sufficient numbers of studies to conduct a separate ALE analysis (although with a stricter cluster criterion). For the other paradigms we examined the results in a qualitative manner with the caveat that the robustness of the respective activation patters has yet to be confirmed. Figure 3A shows significant overlap for studies using the landmark task. The resulting cortical network included no activation of right FEF and DLPF cortex and left aIns, in part due to our more conservative criterion for cluster-level FWE. Otherwise, the pattern was quite similar to that found with the omnibus analysis. While that similarity was expected given that the landmark experiments constituted the largest portion of the omnibus analysis, what was important, the landmark network was remarkably similar to the activation pattern for the grating-scales task, i.e., for the only other existing study that used a purely perceptual paradigm (Chen et al., 2020; Fig. 3B).

**Figure 3.**
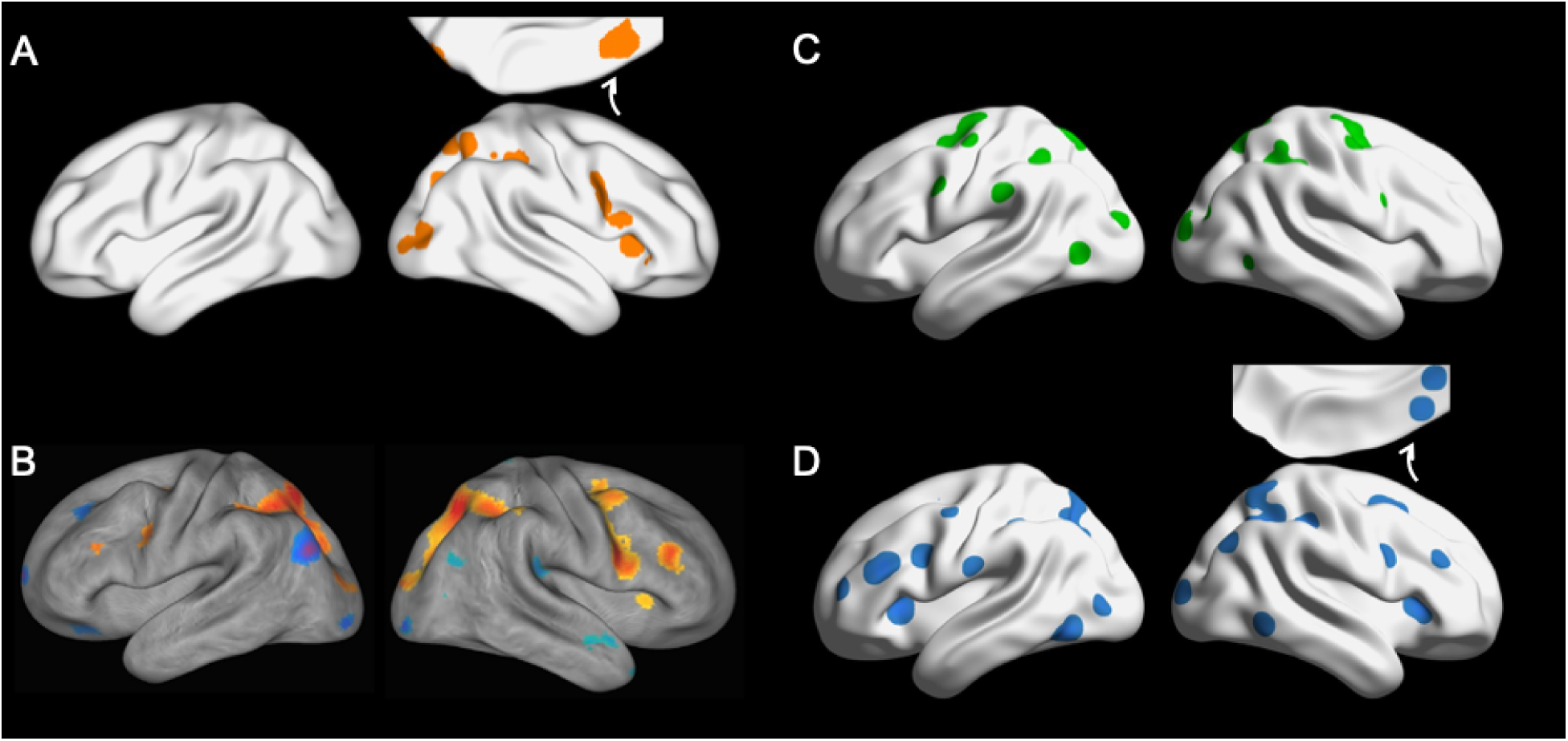
Task-specific analyses. A. Meta-analysis of the landmark task data. B. Partial-least square analysis of the grating-scales task (Chen et al., 2020). C. Overlay plot of the line bisection foci (cerebellar foci not shown). D. Overlay plot of the mental number line foci (calcarine focus not shown).

In addition, to qualitatively examine the influence of action vs. imagery we plotted the foci reported in the six line bisection studies (Fig.3C) and in the two mental number line studies (Fig.3D), respectively. This suggested more bilateral activation with relatively more overlap in the FEFs for manual line bisection as well as more overlap in SPL and DLPFC for mental number lines.

## Discussion

With the current meta-analysis, we intended to investigate the intact functional anatomy underlying perceptual judgment tasks that have been found to give rise to pseudoneglect. We found that in our omnibus analysis across several paradigms (landmark task, line bisection, grating-scales task, and number line task) significant right hemisphere-biased overlap emerged for posterior regions: the IPS, the SPL, and the IOS. Furthermore, we observed right-biased overlap in the FEF, preSMA, DLPFC, caudal IFC and the aIns of the right hemisphere. As we will argue these results add to our understanding of right-dominant functions of perceptual judgments that give rise to perceptual pseudoneglect as well as subtypes of spatial neglect.

However, as one caveat it should be noted that the present meta-analysis might be imperfect. Specifically, the study included far more studies using the landmark task than other paradigms. Such a bias in existing data could result in imbalanced results that would be avoided in a systematically planned conjunction analysis. Despite this imbalance we found that results for the latter type of studies alone were in good agreement with results from a multivariate study using the grating-scales task (Chen et al., 2020; Fig. 3A vs. B).

This consistency also argues against the possibility that the present results merely reflected unspecific activation of the “task-positive” network (e.g., Fox et al., 2005). That is, the pattern of activity reported by Chen and colleagues (2020) was associated with perceptual judgments about the high spatial frequency components of their stimuli relative to a very similar task pertaining to low spatial frequencies. In addition, the present results show a clear asymmetry that was not reported for the task-positive network. In sum, we argue that the current results do serve as a reasonably accurate approximation of the true functional network of brain areas associated with perceptual pseudoneglect tasks. Therefore, in the following paragraphs we will proceed to comparing the current results with those of other methods to then propose a new framework of the neuro-computations underlying perceptual pseudoneglect.

The results of our meta-analysis are partially consistent with the lesion patterns in stroke patients exhibiting line bisection deficits, i.e., lesion sites in the IPS and IOS, medial occipital cortex, and the MFG as reported by Molenberghs et al. (2012).

In contrast to Molenberghs et al. (2012), the current study found the SPL, IFC and anterior insula to be associated with pseudoneglect tasks. This could indicate that these areas merely coactivate in imaging studies without playing a critical role in perceptual judgments. On the other hand, it is possible that lesion studies overlook critical areas, either because these studies are administratively challenging and, thus, often have to deal with methodological imperfections (Moore et al., 2023), or because areas that are critically involved in perceptual judgments in the intact brain are rapidly substituted after lesions (e.g., Pascual-Leone et al., 2005). E.g., some evidence suggests that such neural plasticity starts to play out within minutes of a brain damage (Vesia et al., 2015) which would be outside the temporal scope of lesion studies, potentially resulting in false negative errors.

In sum, we can argue that the current findings offer a largely valid view of the areas that are critically involved in perceptual judgments. Furthermore, this activation pattern can be approximately matched to source localization results from an electroencephalography (EEG) study: Le et al. (2015) observed that activation differences between grating-scales conditions that either do or do not capture pseudoneglect first emerge within roughly bilateral parieto-occipital cortex starting some 74 to 148 ms after stimulus onset and again from 210 to 242 ms, stretching across late stages of the visually evoked P1 component and past the P2, respectively. These sources of activation might be associated with IOS in both hemispheres for the grating-scales task (Fig. 3B) but could involve other visual areas in other tasks (e.g., lateral occipital cortex in the landmark task, Fig. 3A). From 242 to 280 ms then Le et al. (2015) observed clearly right lateralized activation over right posterior cortex (n.b., Le et al., 2015, located the source in the temporo-parietal junction, probably due to imprecise EEG source localization). Based on the current results we believe that this signal originates from right IPS and SPL. From 280 to 340 ms, the same regions are co-actived along with additional sources in interior frontal regions, likely the IFC and aIns. After 340ms, activation becomes confined to those frontal regions, persisting until 394 ms (Le et al., 2015; for the role of the second component of the superior longitudinal fasciculus and its parieto-frontal connectivity during the line bisection task also see Thiebaut de Schotten et al., 2005; Vallar et al., 2014).

This putative temporo-spatial pattern suggests a network that first activates posterior portions of the dorsal attention network before involving anterior portions of the ventral attentional network (Corbetta & Shulman, 2002; Vossel et al., 2014). As such it could be viewed as consistent with the activation-orientation hypothesis of pseudoneglect (Bultitude & Aimola Davies, 2006; Reuter-Lorenz et al., 1990; see Anderson, 1996, for a different account of an underlying attentional bias). According to this hypothesis, right hemisphere dominance in perceptual judgment tasks leads to increased activation of the right brain, which in turn causes attention to orient to the left visual field (although see Chen et al. 2014; Chen et al. 2019; Karnath et al., 1998). Nevertheless, it remains unclear what this right-hemisphere dominance entails and why leftward oriented attention leads to biased perceptual judgments. Here, we suggest a more detailed framework informed by the current results.

### A framework for perceptual judgments and pseudoneglect

Considering the functional network observed in the current study together with the EEG data obtained for the grating-scales task (Le et al., 2015) we tentatively propose the following neuro-computational steps that might underlie perceptual pseudoneglect. What is crucial in this context, we argue that these steps should follow a process of perceptual decision making given the fact that perceptual pseudoneglect is based on perceptual judgments:

**1. Visual analysis.** Perceptual judgment tasks trigger visual analysis of the stimulus material in roughly bilaterally activated striate and extrastriate areas.

**2. Integration and magnitude estimation.** Next, retinotopic areas in IPS and SPL activate, predominantly in the right hemisphere, possibly following interhemispheric competition (Le et al., 2015). Notably, the right IPS and SPL respond not only to the contralateral visual half field but also to extensive portions of the ipsilateral half field (Sheremata & Silver, 2015). Thus, they are well-suited to forming integrated representations of the stimulus as a whole, preserving its topography while accounting for the imperfect isometry between the left and right visual field.

**2a. Feedback.** It appears reasonable to assume that these parietal representations activate striate and extrastriate areas, utilizing their high spatial resolution (e.g., Williams et al., 2008; Chambers et al., 2013; Monaco et al., 2017).

**3. Magnitude comparison.** The integrated stimulus representation is then used to compute magnitudes of the task-relevant feature of the stimulus (length of landmark stimuli, luminance of greyscales, spatial frequency of grating-scales etc.; Walsh, 2003) and to compare these magnitudes across the stimulus’ horizontal extent within the right hemisphere’s IPS and SPL.

**4. Perceptual decision and response implementation.** Finally, a perceptual decision regarding the left vs. the right side of the stimulus is made, involving the aIns, as the IFC and insula are associated with forming an action-oriented representation in preparation of a response (e.g., Demanet et al., 2016; Myers et al., 2017). This decision also accounts for activity in the preSMA, which is known to play a role in action selection (e.g., Coull et al., 2016; Mueller et al., 2007), as well as in the FEF, which is associated with shifting attention (e.g., Corbetta & Shulman, 2002).

Although this framework is provisional and requires further investigation, it aligns with perceptual judgment processes. For that reason, we argue that it can explain several key aspects of perceptual pseudoneglect. For instance, stimulus analysis in visual areas implicated in perceptual pseudoneglect (step 1) is consistent with recent finding that visual stimuli impoverished by pixel noise lead to stronger biases in various perceptual judgment tasks (Chen et al., 2019; Chen & Niemeier, 2014). This may be due to stronger activation of earlier visual, in particular parvocellular processes (Niemeier et al., 2008), at the expense of higher-order visual information (e.g., Kayser et al., 2003; Murray et al., 2002). Next, we argue that the formation of integrated representations of visual stimuli in right parietal cortex (step 2) is the primary cause of right-biased brain activation in perceptual judgment tasks. We further argue that within these areas, a more detailed or salient representation of the contralateral visual field leads to right-biased behavioural biases (step 3). The fact that these parietal areas are also involved in attentional cueing paradigms (Corbetta & Shulman, 2002) further explains why attentional cues modulate perceptual pseudoneglect (Bultitude et al., 2006; McCourt et al., 2005; Singh et al., 2011; Toba et al., 2011). Finally, feedback as suggested in step 2a might also play a role in judgment tasks that involve imagery such as mental number line tasks.

In conclusion, in the present study we used a meta-analysis to identify the functional network involved in perceptual pseudoneglect to reduce the risk of false positive and false negative errors of previous imaging studies. Based on the resulting set of brain areas matched against EEG data for their temporal activation pattern, we propose a new framework of neuro-computational steps that explain pseudoneglect. This framework suggests that pseudoneglect arises interhemispheric integration during perceptual judgments, which leads to asymmetric brain activation and biased attention, rather than attentional biases causing asymmetric judgments. This framework and intact functional anatomy of perceptual pseudoneglect sheds new light on perceptual forms of spatial neglect after brain damage.

## Acknowledgement

We thank Kathleen Pirog Revill for making the imaging data available to be included in this study.

## References

Anderson B (1996). A mathematical model of line bisection behaviour in neglect. Brain 119(Pt 3):841–850. doi:10.1093/brain/119.3.841

Badzakova-Trajkov G, Häberling IS, Corballis MC (2011) Magical ideation, creativity, handedness, and cerebral asymmetries: a combined behavioural and fMRI study. Neuropsychologia 49(10):2896–2903. 10.1016/j.neuropsychologia.2011.06.016

Badzakova-Trajkov G, Häberling IS, Roberts RP, Corballis MC (2010) Cerebral asymmetries: complementary and independent processes. PLoS One 5(3):e9682. 10.1371/journal.pone.0009682

Bates E, Wilson SM, Saygin AP, Dick F, Sereno MI, Knight RT, Dronkers NF (2003) Voxel-based lesion-symptom mapping. Nat Neurosci 6(5):448–450. 10.1038/nn1050

Benwell CS, Harvey M, Thut G (2014) On the neural origin of pseudoneglect: EEG-correlates of shifts in line bisection performance with manipulation of line length. Neuroimage 86(100):370–380. 10.1016/j.neuroimage.2013.10.014

Bowers D, Heilman KM (1980) Pseudoneglect: effects of hemispace on a tactile line bisection task. Neuropsychologia 18(4-5):491–498. 10.1016/0028-3932(80)90151-7

Bultitude JH, Aimola Davies AM (2006) Putting attention on the line: investigating the activation-orientation hypothesis of pseudoneglect. Neuropsychologia 44(10):1849–1858. 10.1016/j.neuropsychologia.2006.03.001

Cai Q, Van der Haegen L, Brysbaert M (2013) Complementary hemispheric specialization for language production and visuospatial attention. Proc Natl Acad Sci U S A 110(4):E322–E330. 10.1073/pnas.1212956110

Caramazza A (1984) The logic of neuropsychological research and the problem of patient classification in aphasia. Brain Lang 21(1):9–20. 10.1016/0093-934x(84)90032-4

Cavézian C, Valadao D, Hurwitz M, Saoud M, Danckert J (2012) Finding centre: ocular and fMRI investigations of bisection and landmark task performance. Brain Res 1437:89–103. 10.1016/j.brainres.2011.12.002

Chambers CD, Allen CP, Maizey L, Williams M A (2013) Is delayed foveal feedback critical for extra-foveal perception? Cortex 49(1):327–335. 10.1016/j.cortex.2012.03.007

Chen J, Kaur J, Abbas H, Wu M, Luo W, Osman S, Niemeier M (2019) Evidence for a common mechanism of spatial attention and visual awareness: Towards construct validity of pseudoneglect. PLoS One 14(3):e0212998. 10.1371/journal.pone.0212998

Chen J, Lee ACH, O’Neil EB, Abdul-Nabi M, Niemeier M (2020) Mapping the anatomy of perceptual pseudoneglect: A multivariate approach. Neuroimage 207:116402. 10.1016/j.neuroimage.2019.116402

Chen J, Niemeier M (2014) Distractor removal amplifies spatial frequency-specific crossover of the attentional bias: A psychophysical and Monte Carlo simulation study. Exp Brain Res 232(12):4001–4019. 10.1007/s00221-014-4082-y

Ciçek M, Deouell LY, Knight RT (2009) Brain activity during landmark and line bisection tasks. Front Hum Neurosci 3:7. 10.3389/neuro.09.007.2009

Corbetta M, Shulman GL (2011) Spatial neglect and attention networks. Annu Rev Neurosci 34:569–599. 10.1146/annurev-neuro-061010-113731

Corbetta M, Shulman GL (2002) Control of goal-directed and stimulus-driven attention in the brain. Nat Rev Neurosci 3(3):201–215. 10.1038/nrn755

Coull JT, Vidal F, Burle B (2016). When to act, or not to act: that’s the SMA’s question. Curr Opin Behav Scie 8:14–21. 10.1016/j.cobeha.2016.01.003

de Haan B, Karnath HO (2018) A hitchhiker’s guide to lesion-behaviour mapping. Neuropsychologia 115:5–16. 10.1016/j.neuropsychologia.2017.10.021

Demanet J, Liefooghe B, Hartstra E, Wenke D, De Houwer J, Brass M (2016) There is more into ‘doing’ than ‘knowing’: The function of the right inferior frontal sulcus is specific for implementing versus memorising verbal instructions. Neuroimage 141:350–356. 10.1016/j.neuroimage.2016.07.059

Eickhoff SB, Bzdok D, Laird AR, Kurth F, Fox PT (2012). Activation likelihood estimation meta-analysis revisited. Neuroimage 59(3):2349–2361. doi:10.1016/j.neuroimage.2011.09.017

Filimon F, Nelson JD, Huang RS, Sereno MI (2009) Multiple parietal reach regions in humans: Cortical representations for visual and proprioceptive feedback during on-line reaching. J Neurosci 29(9):2961–2971. 10.1523/JNEUROSCI.3211-08.2009

Fink GR, Marshall JC, Shah NJ, Weiss PH, Halligan PW, Grosse-Ruyken M, … Freund HJ (2000) Line bisection judgments implicate right parietal cortex and cerebellum as assessed by fMRI. Neurology 54(6):1324–1331. 10.1212/wnl.54.6.1324

Fink GR, Marshall JC, Weiss PH, Shah NJ, Toni I, Halligan PW, Zilles K (2000) ’Where’ depends on ‘what’: A differential functional anatomy for position discrimination in one-versus two-dimensions. Neuropsychologia 38(13):1741–1748. 10.1016/s0028-3932(00)00078-6

Fink GR, Marshall JC, Weiss PH, Stephan T, Grefkes C, Shah NJ, … Dieterich M (2003) Performing allocentric visuospatial judgments with induced distortion of the egocentric reference frame: An fMRI study with clinical implications. Neuroimage 20(3):1505–1517. 10.1016/j.neuroimage.2003.07.006

Fink GR, Marshall JC, Weiss PH, Toni I, Zilles K (2002) Task instructions influence the cognitive strategies involved in line bisection judgments: Evidence from modulated neural mechanisms revealed by fMRI. Neuropsychologia 40(2):119–130. 10.1016/s0028-3932(01)00087-2

Fink GR, Marshall JC, Weiss PH, Zilles K (2001) The neural basis of vertical and horizontal line bisection judgments: An fMRI study of normal volunteers. Neuroimage 14(1 Pt 2):S59-S67. 10.1006/nimg.2001.0819

Fox MD, Snyder AZ, Vincent JL, Corbetta M, Van Essen DC, Raichle ME (2005). The human brain is intrinsically organized into dynamic, anticorrelated functional networks. Proc Natl Acad Sci U S A 102(27):9673–9678. 10.1073/pnas.0504136102

Flöel A, Jansen A, Deppe M, Kanowski M, Konrad C, Sommer J, Knecht S (2005). Atypical hemispheric dominance for attention: functional MRI topography. J Cereb Blood Flow Metab 25(9):1197–1208. https://oi.org10.1038/sj.jcbfm.9600114

Harvey M, Milner AD, Roberts RC (1995) An investigation of hemispatial neglect using the Landmark Task. Brain Cogn 27(1):59–78. 10.1006/brcg.1995.1004

Heilman, K., Watson, R., Valenstein, E. (1993) Neglect and related disorders. In Heilman K Valenstein E (eds) Clinical Neuropsychology, Oxford University Press, New York, pp 279–336.

Hurwitz M, Valadao D, Danckert J (2011) Functional MRI of dynamic judgments of spatial extent. Exp Brain Res 214(1):61–72. 10.1007/s00221-011-2806-9

Jewell G, McCourt ME (2000) Pseudoneglect: A review and meta-analysis of performance factors in line bisection tasks. Neuropsychologia 38(1):93–110. 10.1016/s0028-3932(99)00045-7

Karnath HO, Niemeier M, Dichgans J (1998) Space exploration in neglect. Brain 121(Pt 12):2357–2367. 10.1093/brain/121.12.2357

Kayser C, Salazar RF, Konig P (2003) Responses to natural scenes in cat V1. J Neurophysiol 90(3):1910–1920. 10.1152/jn.00195.2003

Koch G, Oliveri M, Cheeran B, Ruge D, Lo Gerfo E, Salerno S, … Caltagirone C (2008) Hyperexcitability of parietal-motor functional connections in the intact left-hemisphere of patients with neglect. Brain 131(Pt 12):3147–3155. 10.1093/brain/awn273

Le A, Stojanoski BB, Khan S, Keough M, Niemeier M (2015) A toggle switch of visual awareness? Cortex 64:169–178. 10.1016/j.cortex.2014.09.015

Liu D, Zhou D, Li M, Li M, Dong W, Verguts T, Chen Q (2019) The Neural Mechanism of Number Line Bisection: A fMRI study. Neuropsychologia 129:37–46. 10.1016/j.neuropsychologia.2019.03.007

Marshall RS, Lazar RM, Van Heertum, RL, Esser PD, Perera GM, Mohr JP (1997) Changes in regional cerebral blood flow related to line bisection discrimination and visual attention using HMPAO-SPECT. Neuroimage 6(2):139–144. 10.1006/nimg.1997.0283

Mattingley JB, Berberovic N, Corben L, Slavin MJ, Nicholls ME, Bradshaw JL (2004) The greyscales task: a perceptual measure of attentional bias following unilateral hemispheric damage. Neuropsychologia 42(3):387–394. 10.1016/j.neuropsychologia.2003.07.007

McCourt ME, Garlinghouse M, Reuter-Lorenz PA (2005) Unilateral visual cueing and asymmetric line geometry share a common attentional origin in the modulation of pseudoneglect. Cortex 41(4):499–511. 10.1016/s0010-9452(08)70190-4

McCourt ME, Jewell G (1999) Visuospatial attention in line bisection: stimulus modulation of pseudoneglect. Neuropsychologia 37(7):843–855. 10.1016/s0028-3932(98)00140-7

Molenberghs P, Sale MV, Mattingley JB (2012). Is there a critical lesion site for unilateral spatial neglect? A meta-analysis using activation likelihood estimation. Front Hum Neurosci 6:78. 10.3389/fnhum.2012.00078

Monaco S, Gallivan JP, Figley TD, Singhal A, Culham JC (2017) Recruitment of Foveal Retinotopic Cortex During Haptic Exploration of Shapes and Actions in the Dark. J Neurosci, 37(48):11572–11591. 10.1523/JNEUROSCI.2428-16.2017

Moore MJ, Milosevich E, Mattingley JB, Demeyere N (2023) The neuroanatomy of visuospatial neglect: A systematic review and analysis of lesion-mapping methodology. Neuropsychologia 180:108470. 10.1016/j.neuropsychologia.2023.108470

Mueller VA, Brass M, Waszak F, Prinz W (2007). The role of the preSMA and the rostral cingulate zone in internally selected actions. Neuroimage 37(4):1354–1361. 10.1016/j.neuroimage.2007.06.018

Müller VI, Cieslik EC, Laird AR, Fox PT, Radua J, Mataix-Cols D, … Eickhoff SB (2018) Ten simple rules for neuroimaging meta-analysis. Neurosci Biobehav Rev 84:151–161. 10.1016/j.neubiorev.2017.11.012

Murray SO, Kersten D, Olshausen BA, Schrater P, Woods DL (2002) Shape perception reduces activity in human primary visual cortex. Proc Natl Acad Sci U S A99(23):15164–15169. 10.1073/pnas.192579399

Myers NE, Stokes MG, Nobre AC (2017) Prioritizing Information during Working Memory: Beyond Sustained Internal Attention. Trends Cogn Sci 21(6):449–461. 10.1016/j.tics.2017.03.010

Niemeier M, Singh VV, Keough M, Akbar N (2008) The perceptual consequences of the attentional bias: evidence for distractor removal. Exp Brain Res 189(4):411–420. 10.1007/s00221-008-1438-1

Niemeier M, Stojanoski B, Greco AL (2007) Influences of time and spatial frequency on the perceptual bias: evidence for competition between hemispheres. Neuropsychologia 45(5):1029–1040. 10.1016/j.neuropsychologia.2006.09.006

Pascual-Leone A, Amedi A, Fregni F, Merabet LB (2005) The plastic human brain cortex. Annu Rev Neurosci 28:377–401. 10.1146/annurev.neuro.27.070203.144216

Plewan T, Weidner R, Eickhoff SB, Fink GR (2012) Ventral and dorsal stream interactions during the perception of the Müller-Lyer illusion: evidence derived from fMRI and dynamic causal modeling. J Cogn Neurosci 24(10):2015–2029. 10.1162/jocn_a_00258

Price CJ, Friston KJ (1997) Cognitive conjunction: a new approach to brain activation experiments. Neuroimage 5(4 Pt 1):261-270. 10.1006/nimg.1997.0269

Revill KP, Karnath HO, Rorden C (2011) Distinct anatomy for visual search and bisection: a neuroimaging study. Neuroimage 57(2):476–481. 10.1016/j.neuroimage.2011.04.066

Sahan MI, Majerus S, Andres M, Fias W (2019) Functionally distinct contributions of parietal cortex to a numerical landmark task: An fMRI study. Cortex 114:28–40. 10.1016/j.cortex.2018.11.005

Schuster V, Herholz P, Zimmermann KM, Westermann S, Frässle S, Jansen A (2017) Comparison of fMRI paradigms assessing visuospatial processing: Robustness and reproducibility. PLoS One 12(10):e0186344. 10.1371/journal.pone.0186344

Sheremata SL, Silver MA (2015) Hemisphere-dependent attentional modulation of human parietal visual field representations. J Neurosci 35(2):508–517. 10.1523/JNEUROSCI.2378-14.2015

Singh VVW, Stojanoski B, Le A, Niemeier M (2011) Spatial frequency-specific effects on the attentional bias: Evidence for two attentional systems. Cortex 47(5):547–556. 10.1016/j.cortex.2010.03.006

Sperber C, Karnath HO (2017) Impact of correction factors in human brain lesion-behavior inference. Hum Brain Mapp 38(3):1692–1701. 10.1002/hbm.23490

Szczepanski SM, Kastner S (2013) Shifting attentional priorities: Control of spatial attention through hemispheric competition. J Neurosci 33(12):5411–5421. 10.1523/JNEUROSCI.4089-12.2013

Thiebaut de Schotten M, Urbanski M, Duffau H, Volle E, Lévy R, Dubois B, Bartolomeo P (2005). Direct evidence for a parietal-frontal pathway subserving spatial awareness in humans. Science 309(5744):2226–2228. doi:10.1126/science.1116251

Thiebaut de Schotten M, Tomaiuolo F, Aiello M, Merola S, Silvetti M, Lecce F, … Doricchi F (2014) Damage to white matter pathways in subacute and chronic spatial neglect: a group study and 2 single-case studies with complete virtual “in vivo” tractography dissection. Cereb Cortex 24(3):691–706. 10.1093/cercor/bhs351

Toba MN, Cavanagh P, Bartolomeo P (2011). Attention biases the perceived midpoint of horizontal lines. Neuropsychologia 49(2):238–246. doi:10.1016/j.neuropsychologia.2010.11.022

Vallar G, Bello L, Bricolo E, Castellano A, Casarotti A, Falini A, Riva M, Fava E, Papagno C (2014). Cerebral correlates of visuospatial neglect: a direct cerebral stimulation study. Hum Brain Mapp 35(4):1334–1350. doi:10.1002/hbm.22257

Vesia M, Niemeier M, Black SE, Staines WR (2015) The time course for visual extinction after a ‘virtual’ lesion of right posterior parietal cortex. Brain Cogn 98:27–34. 10.1016/j.bandc.2015.05.003

Vossel S, Geng JJ, Fink GR (2014) Dorsal and ventral attention systems: distinct neural circuits but collaborative roles. Neuroscientist 20(2):150–159. 10.1177/1073858413494269

Walsh V (2003) A theory of magnitude: Common cortical metrics of time, space and quantity. Trends Cogn Sci 7(11):483–488. 10.1016/j.tics.2003.09.002

Weidner R, Fink GR (2007) The neural mechanisms underlying the Müller-Lyer illusion and its interaction with visuospatial judgments. Cereb Cortex 17(4):878–884. 10.1093/cercor/bhk042

Weiss PH, Marshall JC, Wunderlich G, Tellmann L, Halligan PW, Freund HJ, … Fink GR (2000) Neural consequences of acting in near versus far space: a physiological basis for clinical dissociations. Brain 123(Pt 12):2531–2541. 10.1093/brain/123.12.2531

Weiss PH, Marshall JC, Zilles K, Fink, GR (2003) Are action and perception in near and far space additive or interactive factors? Neuroimage 18(4):837–846. 10.1016/s1053-8119(03)00018-1

Williams MA, Baker CI, Op de Beeck, HP, Shim WM, Dang S, Triantafyllou C, Kanwisher N (2008) Feedback of visual object information to foveal retinotopic cortex. Nat Neurosci 11(12):1439–1445. 10.1038/nn.2218

Zago L, Petit L, Jobard G, Hay J, Mazoyer B, Tzourio-Mazoyer N, … Mellet E (2017) Pseudoneglect in line bisection judgement is associated with a modulation of right hemispheric spatial attention dominance in right-handers. Neuropsychologia 94:75–83. 10.1016/j.neuropsychologia.2016.11.024

Zago L, Petit L, Mellet E, Jobard G, Crivello F, Joliot M, … Tzourio-Mazoyer N (2016) The association between hemispheric specialization for language production and for spatial attention depends on left-hand preference strength. Neuropsychologia 93(Pt B):394–406. 10.1016/j.neuropsychologia.2015.11.018

